# Presence of vitamin B_12_ metabolism in the last common ancestor of land plants

**DOI:** 10.1101/2023.06.07.544158

**Authors:** Richard G. Dorrell, Charlotte Nef, Setsen Altan-Ochir, Chris Bowler, Alison G. Smith

## Abstract

Vitamin B_12_, or cobalamin, (hereinafter B_12_) is an essential organic micronutrient, required by humans as a cofactor for methionine synthase (METH) and for methylmalonyl CoA mutase (MCM), involved in the propionate shunt. B_12_ is a complex corrinoid molecule made only by a subset of bacteria. Plants and fungi have an alternative methionine synthase (METE) that does not need a B_12_ cofactor, so these organisms are typically considered to neither synthesise nor utilise B_12_. In contrast many algal species utilise B_12_ if it is available, because they encode both METE and METH. Moreover, a large proportion of algal species encode METH only, and so are like animals in being dependent on an external source of the vitamin. Here, we performed a detailed phylogenetic analysis of the distribution of METE, METH and eleven further proteins implicated in B_12_ metabolism in eukaryotic cells across an exhaustive library of over 1,500 plant and algal genomes and transcriptomes. The results reveal the hitherto undetected existence of B_12_-associated metabolism deep into the streptophytes. The B_12_-dependent synthase METH, and the accessory proteins MTRR, CblB, CblC, CblD and CblJ were detected in the basally divergent plant lineage of hornworts, and CblB and CblJ were further identified in liverworts. Using phylogenetic and PFAM analysis we demonstrate this is due to retention of ancestral B_12_-metabolism pathways in the last common ancestor of land plants, followed by at least two independent complete losses in mosses and vascular plants. We further show more limited distributions of genes encoding B_12_-related proteins across the algal tree of life, including MCM and type II ribonucleotide reductase, alongside an obligate B_12_-dependency across several major marine algal orders. Finally, by considering the functional biology of early-diverging land plants, together with the collection sites of ten further algal species inferred to have lost B_12_-dependent metabolism, we propose freshwater-to-land transitions and symbiotic associations to have been major constraining factors in B_12_ availability in early plant evolution.

## Introduction

Vitamins are organic micronutrients that are taken up by organisms from the environment and are essential for central metabolic processes [1, 2]. In particular, B-vitamins are the precursors to enzyme cofactors, providing additional chemical reactivity to that of amino acids. Plants, fungi and microorganisms also require these compounds for their metabolism, but are able to synthesise them *de novo*. The exception is for vitamin B_12_, a cobalt-containing corrinoid molecule that is not synthesised by any eukaryote [3, 4]. In humans, B_12_ is essential for two enzymatic activities: B_12_-dependent methionine synthase (METH) in the C1 cycle [5, 6] and methylmalonyl-coA mutase (MCM), responsible for the metabolism of odd-chain fatty acids and branched-chain amino acids via the propionate shunt [7, 8]. B_12_ deficiency and its associated pathologies (pernicious anaemia, methylmalonic acidemia) may be chronic in subsistence economies, with particularly adverse impacts on child development and during pregnancy [9–11]. Addressing how to mitigate B_12_ deficiency in plant-based diets, which may have otherwise substantially lower environmental impacts than omnivorous nutrition, is therefore a key challenge to sustainably feeding a growing planetary population [12–14].

Historically, B_12_ in the human diet has been obtained from different sources. These include from ruminant animals and their derivatives (i.e., dairy products) [15], the direct consumption of B_12_ from soil particles (e.g., via geophagia [16], from fortified (nutritional yeast) and from edible seaweeds from across the algal tree of life [15, 17]. Indeed, many microalgae, microscopic photosynthetic eukaryotes, also encode METH and/or MCM and many are obligately dependent on B_12_ [18, 19]. In one study for example, 171/326 sampled algal species were found to be unable to grow in the absence of supplemented B_12_ [18]. B_12_ auxotrophy has been subsequently documented to be widespread in specific algal phyla (e.g., haptophytes, dinoflagellates) across the Tree of Life [20, 21]. Some microalgal lineages may encode further B_12_-dependent enzymes (e.g. form II ribonucleotide reductase (RNRII), first documented in the green microalga *Euglena*) [7], which is not found in humans.

B_12_ biosynthesis involves over twenty discrete enzymatic steps from the common tetrapyrrole intermediate uroporphyrinogen III [22, 23]. To date, complete B_12_ biosynthesis has only been described in a subset of bacteria and archaea [22, 24]. Algae may acquire synthesised B_12_ by scavenging from the environment [25], phagotrophic consumption of B_12_-containing organisms [26] or via symbiotic exchanges with B_12_-producing commensals [18]. Additionally, there are a number of B_12_ variants with different axial ligands, which affects the bioavailability of the vitamin. Intrinsic factor, the B_12_-binding protein found in the human ileum, has a much higher affinity for cobalamin, where the lower axial ligand is 5,6-dimethylbenzimidazole (DMB), than for pseudocobalamin, which has adenine as the lower axial ligand [27]. Eukaryotic microalgae have a similar preference for cobalamin over pseudocobalamin [27], although a B_12_ remodelling pathway has been documented in some species [27, 28], which enables them to convert pseudocobalamin to cobalamin if supplied with DMB.

Little is known of the uptake, cellular transport or metabolism of B_12_ in algae, but the pathways are well documented in humans. Following its endocytic internalisation into the cell, cobalamin is released into the cytosol via the co-operative activity of a cytosolic chaperone CblC and the lysosomal transporters CblF/ CblJ [29] (**Fig. 1**). This involves the reductive removal of any preceding upper-axial ligands associated with the cob(III)-alamin and conversion into cob(I)- and cob(II)alamins complexed with CblC [29]. Cob(I,II)-alamins complexed with CblC can then be transferred to the conjugate protein CblD, which subsequently directs the synthesis of different cobalamin-dependent enzymes [29, 30]. METH requires methyl-cobalamin, produced from CblD-associated B_12_ by the activity of methionine synthase reductase (MTRR) [5] in the cytosol, whereas synthesis of MCM involves adenosyl-cobalamin, produced in the mitochondria via the conjugate activity of the probable B_12_ transporter protein CblA [31] and the adenosyl-transferase CblB [7]. A small number of additional proteins (e.g., CblX, epi-CblC) have been shown to have epistatic effects on cobalamin uptake in humans, but are of unknown function [29]. Finally, in the algal species in which it occurs, RNR-II is cytosolic and utilises adenosyl-cobalamin.

**Fig. 1.**
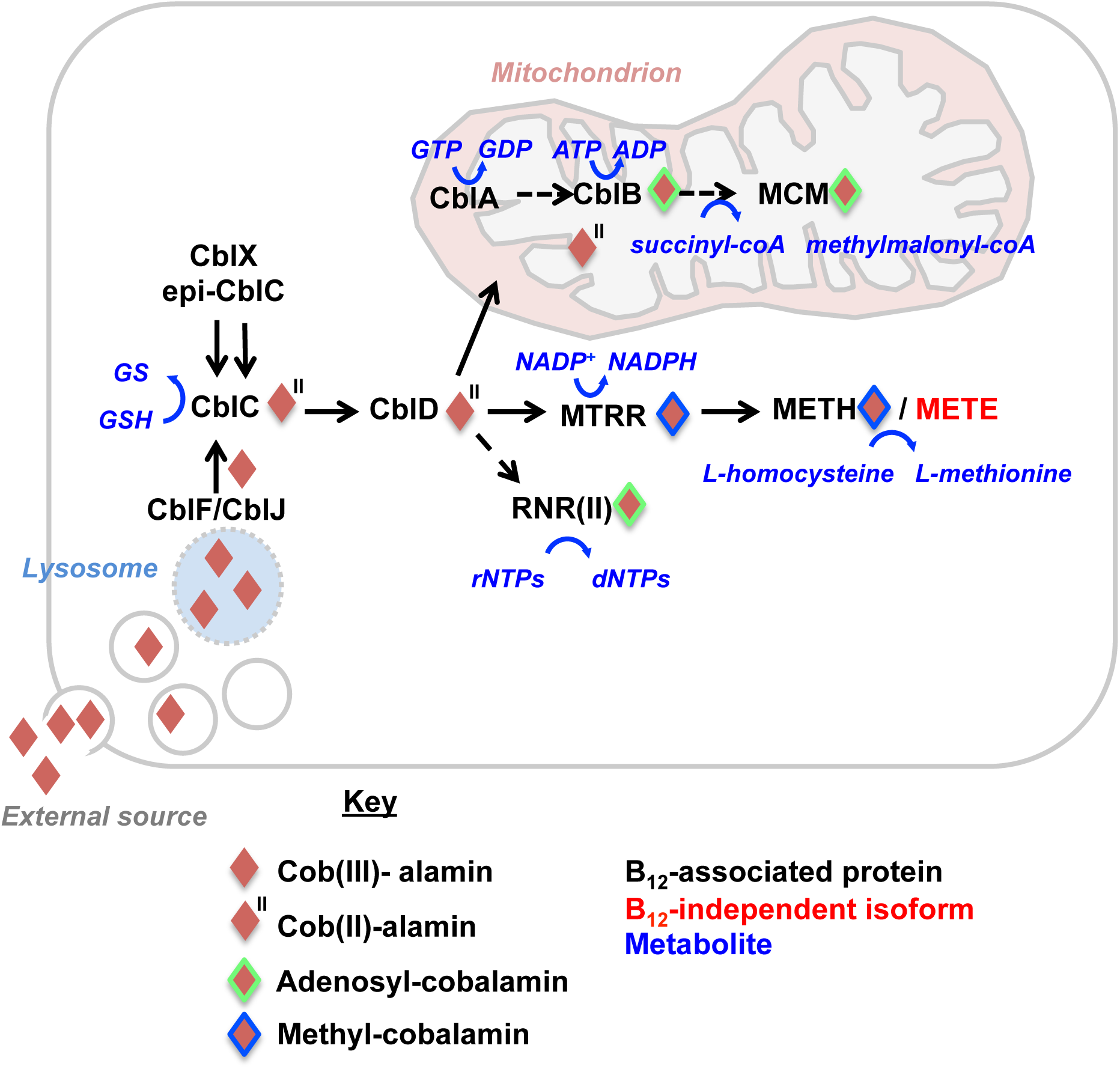
Known B_12_ uptake and utilisation pathways in eukaryotes. This figure shows a schematic eukaryotic cell, with potential B_12_-associated proteins detected in previous studies across the eukaryotes, labelled following [29] and [36]. The geneIDs, origins and PFAM domains associated with each query protein for the phylogenetic analysis are tabulated below.

B_12_ produced by bacteria may be scarce in the environment, including in soil [32] and large portions of the contemporary ocean [33, 34], and uptake and processing of the vitamin is energetically costly [29, 35]. In contrast to the widespread utilisation of B_12_ by aquatic algae, several ecologically successful groups of eukaryotes, including all previously documented land plants, have lost their metabolic dependence on B_12_ [36]. This is possible because of a B_12_-independent methionine synthase (METE) that arose separately, which can replace METH, albeit with a lower catalytic activity [37]. The METE gene shows a discontinuous presence across the tree of life, and is also present in many METH-containing species, including a substantial proportion of eukaryotic algae, which renders them facultative B_12_ users [36]. The most parsimonious explanation is an ancestral presence of both METE and METH genes in eukaryotes, with the repeated loss of METE from obligately B_12_-dependent species, and the occasional loss of METH from plants, fungi, and other species that do not have B_12_-associated metabolism.

In the last five years, the dramatic expansion in plant and algal genomic and transcriptomic resources, such as through the OneKp project [38, 39], have provided unprecedented insights into the foundational events underpinning plant evolution. This includes evidence for the stepwise accumulation of evolutionary innovations (e.g., homeobox domain transcription factors, auxin transport pathways and water-stress tolerance factors) associated with the colonisation of land in the closest streptophyte relatives of land plants [40–42], and the importance of gene losses as well as gene family expansions for the post-terrestrial diversification of both bryophyte (mosses, liverworts, and hornworts) and vascular plants [43, 44]. These expanded genomic resources prompted us to revise our current understanding of the distribution of B_12_-dependent metabolism across the photosynthetic eukaryotic tree of life. Here we investigate the present of B_12_ uptake and associated metabolism in over 1,600 published plant and algal genome and transcriptome libraries, and consider in which ecological contexts loss of B_12_-dependent metabolism has occurred.

## Results

### Distribution of METH/ METE genes indicate B_12_ presence in hornworts

We identified homologues of the thirteen B_12_-associated proteins, illustrated in **Fig. 1**, from the algae *Chlamydomonas reinhardtii, Euglena gracilis* and *Phaeodactylum tricornutum* and from humans (Table 1), and used these to search a composite library of 1,666 plant and algal genomes and transcriptomes. The library contained: 44 algal and plant genomes, including the recently published draft genomes of the hornworts *Anthoceros agrestis* and *A. punctigera* [44, 45]; decontaminated versions of the 1,000 plant transcriptomes (OneKp ; 1,292 libraries) [38, 46] and the Marine Microbial Eukaryote Transcriptome Sequencing Project (MMETSP; 300 libraries) [47, 48]; and 26 further eukaryotic transcriptomes, in particular sampling ecologically abundant marine diatom genera [49] and freshwater chrysophyte lineages [50] (**Table 2**). Homologues were retrieved by reciprocal BLAST best-hit (RbH), single-gene RAxML trees, and PFAM domain analysis (Materials and Methods) [51–53], and were categorised taxonomically following recently published multigene phylogenies of plant and algal diversity [38, 49]. Full outputs are provided in **Dataset S1**.

**Table 1.** Overview of the query sequences used for phylogenetic searches, as obtained from human [29] and algal gene sequences [36].

| **Protein** | **GeneID** | **Query Source** | **PFAMs** |
| --- | --- | --- | --- |
| METH | XP_042923308.1 | *Chlamydomonas* | PF00809, PF02310, PF02574, PF02607, PF02965 |
| METE | AAC49178.1 | *Chlamydomonas* | PF01717, PF08267 |
| MCM | XP_002179504.1 | *Phaeodactylum* | PF01642 |
| RNR | Q2PDF6.1 | *Euglena* | PF02867, PF17975 |
| MTRR | AAF16876.1 | Human | PF00175, PF00258, PF00667 |
| CblA | NP_001362573.1 | Human | PF03308 |
| CblB | NP_443077.1 | Human | PF01923 |
| CblC | NP_056321.2 | Human | PF16690 |
| CblD | NP_056517.1 | Human | PF10229 |
| CblF | NP_060838.3 | Human | PF04791 |
| CblJ | NP_001340531.1 | Human | PF00005 |
| CblX | NP_005325.2 | Human | PF01344, PF07646, PF13415, PF13418, PF13854, PF13964 |
| epi-CblC | NP_001189360.1 | Human | PF00578, PF08534, PF10417 |

**Table 2.**
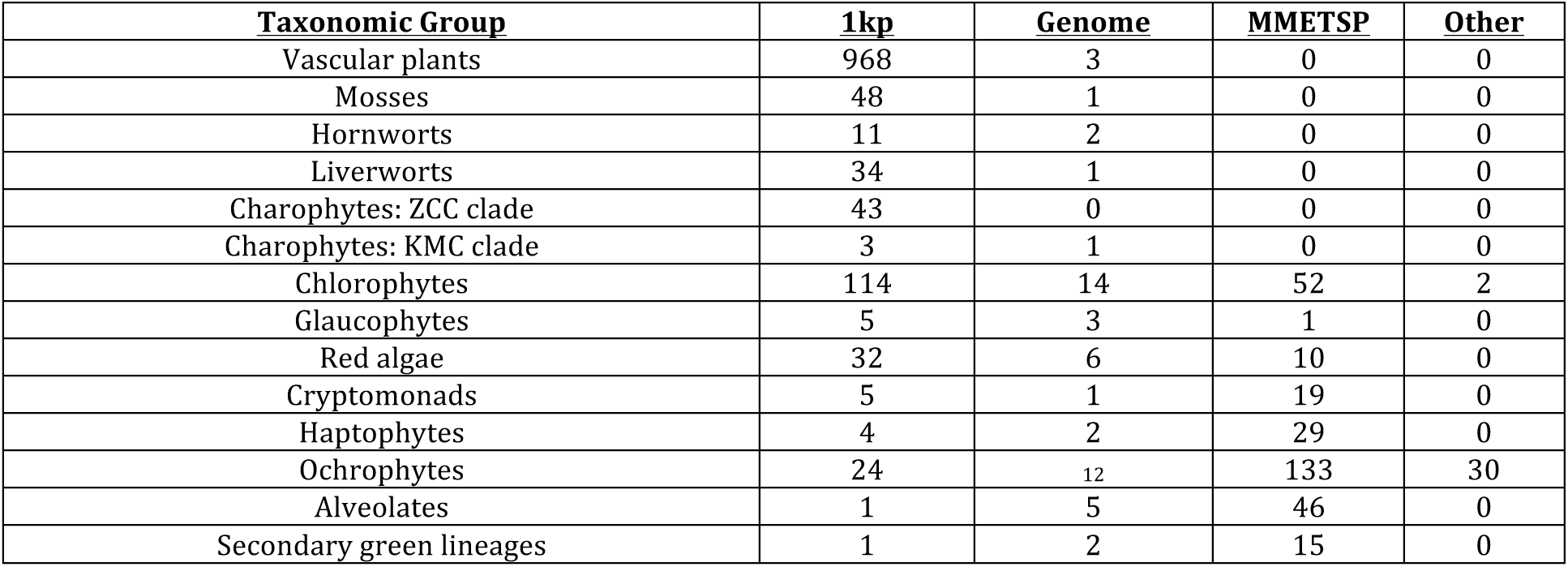
Overview of the libraries searched for presence of B_12_-containing homologues. Totals are provided for the unique numbers of species searched in different taxonomic groups, divided by library type (genome, 1kp transcriptome, MMETSP transcriptome, and other including single-cell genome and independent transcriptome libraries). Totals are provided by species as opposed to strain, and therefore e.g. three searched hornwort genomes (*Anthoceros agrestis* Oxford, *A. agrestis* Bonn, *A. punctigera*) [44] are tabulated as two distinct species. Species for which both genome and 1kp/ MMETSP transcriptomes have been sequenced (e.g., *A. agrestis*) are tabulated in both columns. Acronyms : ZCC-<u>Z</u>ygnematophytes, <u>C</u>oleochaetales, <u>C</u>harophytes ; KMC-<u>K</u>lebsormidiophyes, *<u>M</u>esostigma*, *<u>C</u>hlorokybus*.

| **<u>Taxonomic Group</u>** | **<u>1kp</u>** | **<u>Genome</u>** | **<u>MMETSP</u>** | **<u>Other</u>** |
| --- | --- | --- | --- | --- |
| Vascular plants | 968 | 3 | 0 | 0 |
| Mosses | 48 | 1 | 0 | 0 |
| Hornworts | 11 | 2 | 0 | 0 |
| Liverworts | 34 | 1 | 0 | 0 |
| Charophytes: ZCC clade | 43 | 0 | 0 | 0 |
| Charophytes: KMC clade | 3 | 1 | 0 | 0 |
| Chlorophytes | 114 | 14 | 52 | 2 |
| Glaucophytes | 5 | 3 | 1 | 0 |
| Red algae | 32 | 6 | 10 | 0 |
| Cryptomonads | 5 | 1 | 19 | 0 |
| Haptophytes | 4 | 2 | 29 | 0 |
| Ochrophytes | 24 | 12 | 133 | 30 |
| Alveolates | 1 | 5 | 46 | 0 |
| Secondary green lineages | 1 | 2 | 15 | 0 |

**Table 3.** Identified species inferred to have lost Vitamin B_12_-associated metabolism, based on the retention of METE and minimally the absence of METH and MTRR sequences as verified by phylogenetic and PFAM analysis. A detailed map of the recorded collection sites of all algae included in the dataset is provided at https://tinyurl.com/3esfpkmv.

| Species | Lineage | Strain | Isolation site | Isolation latitude | Isolation longitude | Notes |
| --- | --- | --- | --- | --- | --- | --- |
| Land Plants | Mosses, Vascular plants, liverworts? |  |  |  |  |  |
| *Blastophysa cf. rhizopus* | Chlorophytes | M3368 / BEA 0074B | Epiphyte; Spain, Canary Islands, Fuerteventura, near the lighthouse. Marine | 28,4052029 | -14,4413182 | Based on PFAM presence only |
| *Botryococcus braunii* | Chlorophytes | UTEX 2441 | Lake Huaypo, Peru; Freshwater | -13.5 | -72.1 | Based on PFAM presence only |
| *Brandtodinium nutriculum* | Dinoflagellates | RCC3387 | Marine; foraminiferan endosymbiont | 43.7 | 7.3 |  |
| *Cephaleuros virescens* | Chlorophytes | SAG 28.93 | Natal, parasite from leaves of Camellia | -29.9 | 31.0 | Based on PFAM presence only; Trentepohliales |
| *Coccomyxa pringsheimii* | Chlorophytes | SAG 216.7 | Finland: freshwater; phycobiont of lichen Botrydina vulgaris | N/A | N/A |  |
| *Cyanidioschyzon merolae* | Red algae | 10D | Hot spring; Italy; Sardinia | 40.9 | 14.3 |  |
| *Galdieria sulphuraria* | Red algae | 074W | Fumaroles, Mount Lawu, Indonesia | -7.6 | 11.2 | Encodes MCM, CblA |
| *Geminigera cryophila* | Cryptomonads | CCMP2564 | Marine | -77,8 | -163 |  |
| *Ignatius tetrasporus* | Chlorophytes | UTEX20<sub>12</sub> | Freshwater | N/A | N/A | May encode RNR II |
| *Leptosira obovata* | Chlorophytes | SAG 445.1 | Basel, boggy water at Rosenau; freshwater | 47.5 | 7.9 |  |
| *Porphyridium cruentum* | Red algae | UTEX 161 | Basel, Switzerland; wet shaded tuff | 47.6 | 7.6 | Encodes CblB |
| *Poteriospumella JBC07* | Chrysophytes | JBC07 | Peoples Republic of China, Lake Tai Hu | 31.5 | 120.2 |  |
| *Pseudoscourfieldia marina* | Chlorophytes | SCCAP K/0017 | Inner Oslofjord, Norway | 59.4 | 10.6 | Encodes CblB |
| *Trentepohlia annulata* | Chlorophytes | SAG 20.94 | Damasská cesta pobl. Svratky, Ceskomoravská vrchovina | 49.7 | 16.0 | Based on PFAM presence only; Trentepohliales |
| *Vitrella brassicaformis* | Alveolates | CCMP3155 | Marine, coral | -23.5 | 152 |  |

The phylogenetic distributions of putative homologues are shown schematically in **Fig. 2** for each protein. The patterns underline the widespread occurrence of the B_12_-dependent form of methionine synthase, METH, across eukaryotic algae, found in all major marine groups, and in both early-branching (Klebsormidiaceae, Mesostigmatophyceae, Chlorokybophyceae) and close relatives of land plants (Zygnematophyceae, Charophyceae, Coleochaetophyceae) within the streptophyte lineage [38, 54]. METH was further detected in multiple distantly related hornwort genera, suggesting widespread conservation of B_12_-dependent methionine synthases across the class Anthoceratophyta (**Dataset S1,** sheet 5) [38, 55]. These included OneKp homologues from the hornwort genera *Megaceros* (*M. tosanus*, OneKp transcript-UCRN-2004435; *M. vincentianus*-TCBC-2004163), *Nothoceros* (N. *aenigmaticus* DXOU-2038410), *Paraphymatceros* (*Paraphymato. hallii*: FAJB-2057847) and *Phaeoceros* (*Phaeo. carolinianus*, RXRQ-2022853, RXRQ-2022854), as well as probable METH genes from the *Anthoceros agrestis* Bonn (geneID: Sc2ySwM_228_5027); *A. agrestis* Oxford (geneID: utg000003l_252) and *A. punctigera* (geneID: utg000098l_165) genomes. METH was not found elsewhere within the land plant lineage, except for one potential METH homologue identified by RbH in the liverwort *Blasia sp.* (OneKp transcript: AEXY-2015053; **Dataset S1**, sheet 5), but that resolved during preliminary phylogenetic analyses with bacterial sequences and showed high sequence similarity with a *Mucilaginibacter* METH (NCBId: WP__12_9570389.1; >96% by BLASTp) [56]. Previous studies have documented high levels of bacterial contamination in the *Blasia* transcriptome, [46], and we therefore consider that this is unlikely to correspond to a liverwort METH.

**Fig. 2.**
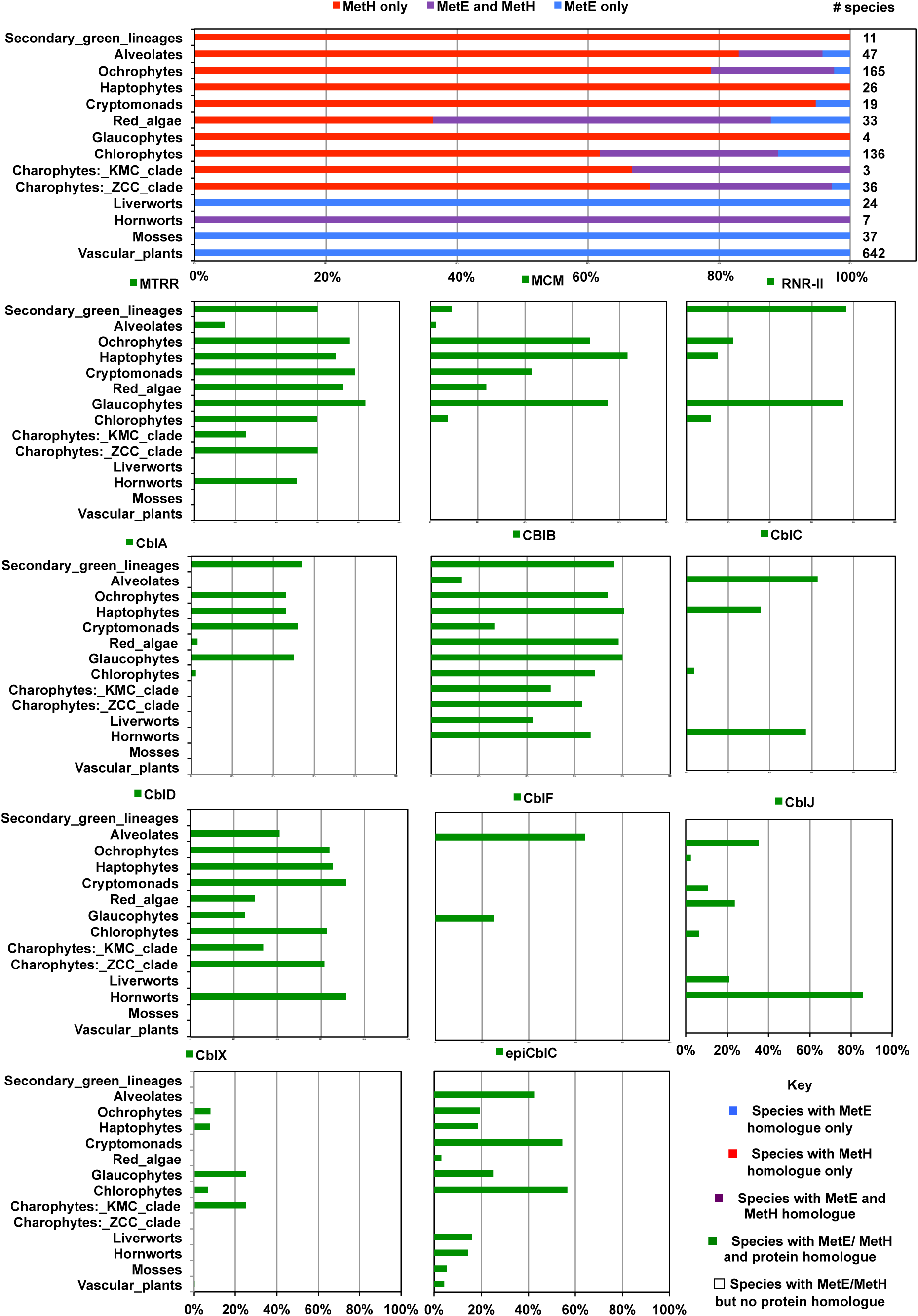
Bar plots of the occurrence of B_12_-associated metabolism across photosynthetic eukaryotes. These graphs show the number of species from thirteen different algal phylogenetic or functional groups inferred to possess METH, METE, or both METH and METE, by combined RbH, PFAM domain and single-gene phylogenetic analysis ; and the number of species from these groups for which at least one B_12_-associated enzyme was detected (the following, plus METE or METH) inferred to possess homologues of MTRR, MCM, RNR II, CBLA, CBLB, CBLC, CBLD, CBLF, CBLJ, CBLX or epi-CBLC via the same methodology. The total number of species assessable in each case is given to the right-hand side of each plot. Abbreviations : KMC-Klebsormidiaceae, Mesostigmatophyceae, Chlorokybophyceae ; ZCC-Zygnematophyceae, Charophyceae, Coleochaetophyceae. Complete tabulated occurrences per lineage and species are provided in **Dataset S1,** sheets 1-2 ; and individual homologue lists per gene in **Dataset S1**, sheets 5-17.

The phylogeny of METH sequences robustly resolved a position for the hornwort sequences within the streptophytes (RAxML bootstrap support : 100%), in a weakly supported sister-group position to homologues from the charophytes / coleochaetales (*Chara vulgaris, Chaetosphaeridium globosum, Coleochaete scutata, Col. irregularis*) (**Fig. 3 ; Dataset S1**). This group was positioned more deeply within Viridiplantae sequences, as a sister to equivalent sequences from Chlorophytes (RAxML bootstrap support: 100% ; **Fig. 3**). Each of the phylogenetically validated *Megaceros, Nothoceros, Phaeoceros* and *Paraphymatoceros* hornwort OneKp sequences possessed all five PFAMs typically associated with METH (homocysteine S-methyltransferase, PF02574 ; pterin-binding, PF00809 ; B_12_-binding, PF02607 and PF02310 ; and the activation domain, PF02965) with similar e-values to functionally characterised equivalents from algae (**Fig. S1**) [21]. The three identified *Anthoceros* homologues possessed all PFAMs apart from the first B_12_-binding (PF02607) domain. However, this PFAM is detected in alternative gene models in each genome (*A. agrestis* Bonn: 362_443; *A. agrestis* Oxford: utg000003l_664; *A. punctigera*, utg000145l_284), suggesting that *Anthoceros* is likely to also possess a functionally active METH. The hornwort sequences all had conserved active sites (e.g., the substrate-binding pocket of the pterin-binding site) associated with METH activity (**Fig. S1**) [21, 57].

**Fig. 3.**
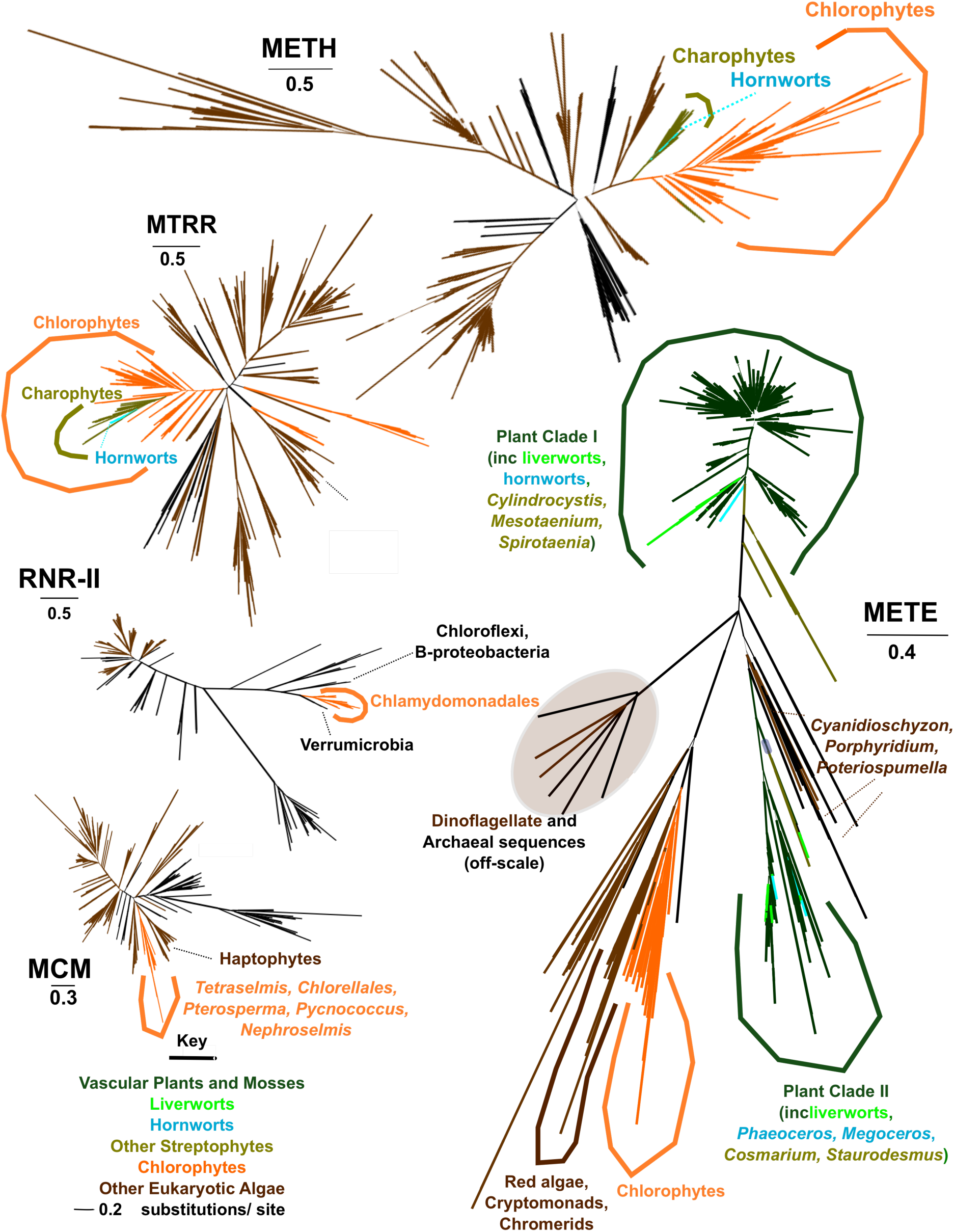
Phylogenetic trees of B_12_-dependent metabolism across photosynthetic eukaryotes. This figure shows best-scoring RAxML trees realised across 350 bootstrap replicates with the PROTGAMMAJTT substitution matrix for METE, METH, MTRR, MCM and RNR II enzymes. Suspected dinoflagellate and archaeal METE sequences are shown in a separate panel to the remaining tree topology, due to the high divergence (> 2.0 substitutions per site) observed from all other aligned sequences. Taxa are coloured by phylogenetic origin: dark green-vascular plants and mosses; light green-liverworts; cyan-hornworts; dark yellow-other streptophytes; orange-chlorophytes; dark brown-other algae; black-other lineages. Branch thickness corresponds to bootstrap support. Initial and curated alignment and tree topologies are provided in **Dataset S1**, sheets 18-19 ; and taxa inferred to be contaminants and thus excluded in **Dataset S1**, sheet 20.

We also considered the distribution of the methionine synthase reductase (MTRR) necessary for METH activity in our dataset. Well-conserved homologues of MTRR were detected across hornwort genomes and transcriptomes, but were absent from other plant groups (**Fig. 2**). As per the situation for METH, the hornwort MTRR genes grouped with other streptophyte sequences within the Viridiplantae (**Fig. 3**), and had well-conserved PFAM domains (flavodoxin 1- PF00258, FAD-binding- PF00667, oxidoreductase- PF00175 ; **Fig. S1**), suggesting vertical origin and functionality. We therefore conclude that the hornwort METH sequences are likely to be catalytically functional, and vertically inherited from the streptophyte ancestor of terrestrial plants.

### Widespread occurrence of two distinct METE isoforms in plants

All plant libraries studied possessed *METE* genes, including hornworts, indicating that both genes were present in the last common plant ancestor (**Fig. 2**). The hornwort *METE* genes resolved with other plant homologues (**Fig. 3**) and possessed well-conserved N-terminal (PF08267) and C-terminal catalytic (PF01717) PFAM domains (**Fig. S1**), indicating their likewise probable vertical origin and functionality.

The plant METE enzymes resolved phylogenetically in two discrete families: a conventional isoform with 1,768 recovered examples (labelled “Clade 1”); and a less frequently observed isoform (“Clade II”) with 125 observed examples (**Fig. 3 ; Dataset S1**). Both isoforms contained credible occurrences of both Meth_synt_1 (PF08267) and Meth_synt_2 (PF01717) domains, and thus are likely to function as methionine synthases (**Fig. S1 ; Dataset S1**). Both Clade I and Clade II isoforms were furthermore detected in higher plant, moss, liverwort, hornwort and streptophyte sequences (**Fig. 3**), suggesting their presence in the last plant common ancestor. All three characterised *Arabidopsis* methionine synthases (ATMS1-At5g17920; ATMS2 At3g03780; ATMS3-At5g20980) resolved within Clade 1, but the Clade II enzymes all showed greater proximity by BLASTp analysis to these than to the four closely related *Arabidopsis* adenosyl-methionine synthases (SAM1-At1g02500; SAM2-At4g01850; SAM3-At3g17390; and SAM4-At2g36880), implying that they likewise correspond to probable methionine synthase isoforms (**Dataset S1**, sheet 6) [58].

Both Clade I and Clade II isoforms distantly positioned in the phylogeny to chlorophyte METE sequences, which formed a sister-group to red algae, cryptomonads and chromerids (**Fig. 3**). The most parsimonious explanation for this distribution would be the ancestral replacement of the streptophyte METE sequence, still retained in chlorophytes, with the Clade I and Clade II isoforms, preceding the loss of METH within specific plant groups.

### Distribution of METE genes confirms widespread B_12_ auxotrophy in algae

In contrast to the widespread distribution of both METE isoforms in land plants, METE was absent from many of the algal libraries searched (**Fig. 2**). Transcriptome libraries may under-report the presence of METE, as it may be transcriptionally repressed in B_12_-supplemented cultures (most common algal growth media include exogenous B_12_, either in the form of soluble vitamins or soil extract) [37, 59], and therefore we limit our consideration to lineages with at least one sequenced genome included [39]. Consistent with previous studies [20], we project B_12_ auxotrophy in all haptophytes (including three genomes included in the study : *Emiliania huxleyi, Chrysochromulina tobinii*, and *Pavlovales* sp. CCMP2436) [60]. Only one potential haptophyte METE homologue (from *Pavlova lutheri* UTEX LB1293) was found by RbH, but was excluded as a probable green algal contaminant (**Dataset S1**). No occurrences of METE were found in the prasinophyte class Mamiellophyceae (including the *Micromonas* and *Ostreococcus* sp. genomes) or the ochrophyte class Pelagophyceae (including two genomes : *Aureococcus anophageferrens,* and Pelagophyceae sp. CCMP2097) [60] (**Fig. 2**). The shared absence of METE from haptophytes, pelagophytes and Mamiellophyceae is particularly interesting given their environmental abundance as assessed e.g., via the *Tara* Oceans expedition [61, 62], underlining the importance of B_12_ acquisition for marine photosynthesis.

Further instances of obligate B_12_ requirements, due to the complete absence of METE, were found in less ecologically abundant algal groups and at more phylogenetically localised scales, including most immediate relatives of the model green algal species *C. reinhardtii* [63, 64] and the glaucophytes and Palmophyllphyceae, respectively considered as basally-divergent members of primary chloroplast-containing algae and the Viridiplantae [65, 66]. Finally, dinoflagellates (including three genomes from *Symbiodinium* sp.) were found to possess only partial METE sequences, which lacked the conventional N-terminal PFAM domain (MetH_synt_1 ; PF08267). The function of this partial METE sequence isoform awaits experimental validation, although B_12_ dependency has been proposed to be widespread across the dinoflagellates [21, 67] (**Dataset S1**). We did, however, identify complete METE genes containing both PFAM domains in the chromerid algae *Chromera velia* and *Vitrella brassicaformis,* which are the closest obligately photosynthetic relatives of dinoflagellates within the alveolates [68]. The dinoflagellate METE-like genes were distantly related to all other equivalents and instead resolved with Archaea (**Fig. 3**), which likewise lacked the N-terminal PFAM domain. This may suggest a horizontal acquisition of the current dinoflagellate METE-like gene that may have been accompanied by the loss of the complete METE isoform.

Finally, the METE genes of the red algal genera *Porphyridium* and *Cyanidioschyzon*, as well as in the chrysophyte *Poteriospumella,* resolved with homologues from bacteria (**Dataset S1** ; **Fig. 3**), independent to all other plant and algal proteins. In each case, these acquisitions were shared between multiple related species or strains, and are therefore likely to be genuine bacterial horizontal gene transfers as opposed to contaminants. Overall, these data indicate complex patterns of METE gene transfer and loss across the photosynthetic eukaryote tree of life.

### MCM and RNR II enzymes are restricted to individual algal lineages

Alongside methionine synthase, we considered the distributions of the B_12_-dependent MCM and RNR II enzymes across our dataset (**Fig. 2**). Neither enzyme was detected in hornworts or more broadly across streptophytes, and were absent from the chlorophytes, except for some sporadic occurrences that are likely consistent with horizontal acquisition as opposed to an ancestral presence in Viridiplantae (discussed below). Homologues of MCM were primarily detected in eukaryotic algae with secondary red chloroplasts (e.g., diatoms. cryptomonads, haptophytes, ochrophytes), where they typically possessed well-conserved PFAM domains, and in many cases predictable mitochondrial targeting sequences consistent with mitochondrial function (**Dataset S1 ; Fig. S2**). MCM is implicated in the catabolism of branched-chain amino acids in diatoms under nitrate exhaustion [69, 70], and has further been proposed to participate in the dissipation of excess diatom plastidial reducing potential [71]. We note that a previous RNAi study of *Phaeodactylum* MCM failed to identify a clear mutant phenotype [70], so it remains to be determined to what extent MCM activity is essential to algae with secondary red chloroplasts, and therefore whether it provides a reason alongside METH for widespread B_12_ dependence across these groups in the environment.

We additionally detected MCM in a small number of glaucophytes, red algae (*Galdieria sulphuraria*) and green algae within the genera *Tetraselmis* and *Chlorella* (**Fig. 2**). In contrast to green algal methionine synthase-associated enzymes, whose phylogenies show typically vertical inheritances, the MCM phylogeny retrieve a sister-group relationship between green algae and haptophytes (**Fig. 3**), implying a probable ancestral loss of the propionate shunt within the Viridiplantae, and subsequent re-acquisition by horizontal gene transfer in specific lineages.

Similarly, the B_12_-dependent type II ribonucleotide reductase (RNR II), first detected in eukaryotes in the secondary green chloroplast-containing alga *Euglena gracilis* [36, 72] was also found in the distantly related haptophytes and chlorarachniophytes, which have secondary green chloroplasts, reflecting possible ancestral horizontal gene transfers inferred between all three lineages [73]. Intriguingly it is also present in the Chlamydomonadales (although not *C. reinhardtii*), where it has most likely been acquired by an independent horizontal gene transfer from bacteria (**Figs. 2, 3**).

### Distribution of uptake proteins suggests deeper retention of B_12_-dependent metabolism within the bryophytes

We further considered the distribution of six proteins associated with the uptake and intracellular transport of B_12_ (CblA, B, C, D, F, and J) and two known epistatic regulators in humans (CblX, epi-CblC) [29] across our dataset. Consistent with the distribution of METH and MTRR, we could identify widespread presence of CblC, D and J proteins in hornwort libraries (**Fig. 2**). The hornwort sequences resolved within clades of other Viridiplantae sequences, and typically contained well-conserved PFAM domains (**Fig. 4** ; **Fig. S1**), suggesting vertical origin and conserved function. The exact location of these enzymes, and precise cellular trafficking pathway in which they may be involved remain to be determined experimentally, as very few carried credible signal peptides that might suggest targeting to the endomembrane system, and participation in a B_12_ endosomal uptake pathway (**Fig. S2**).

**Fig. 4.**
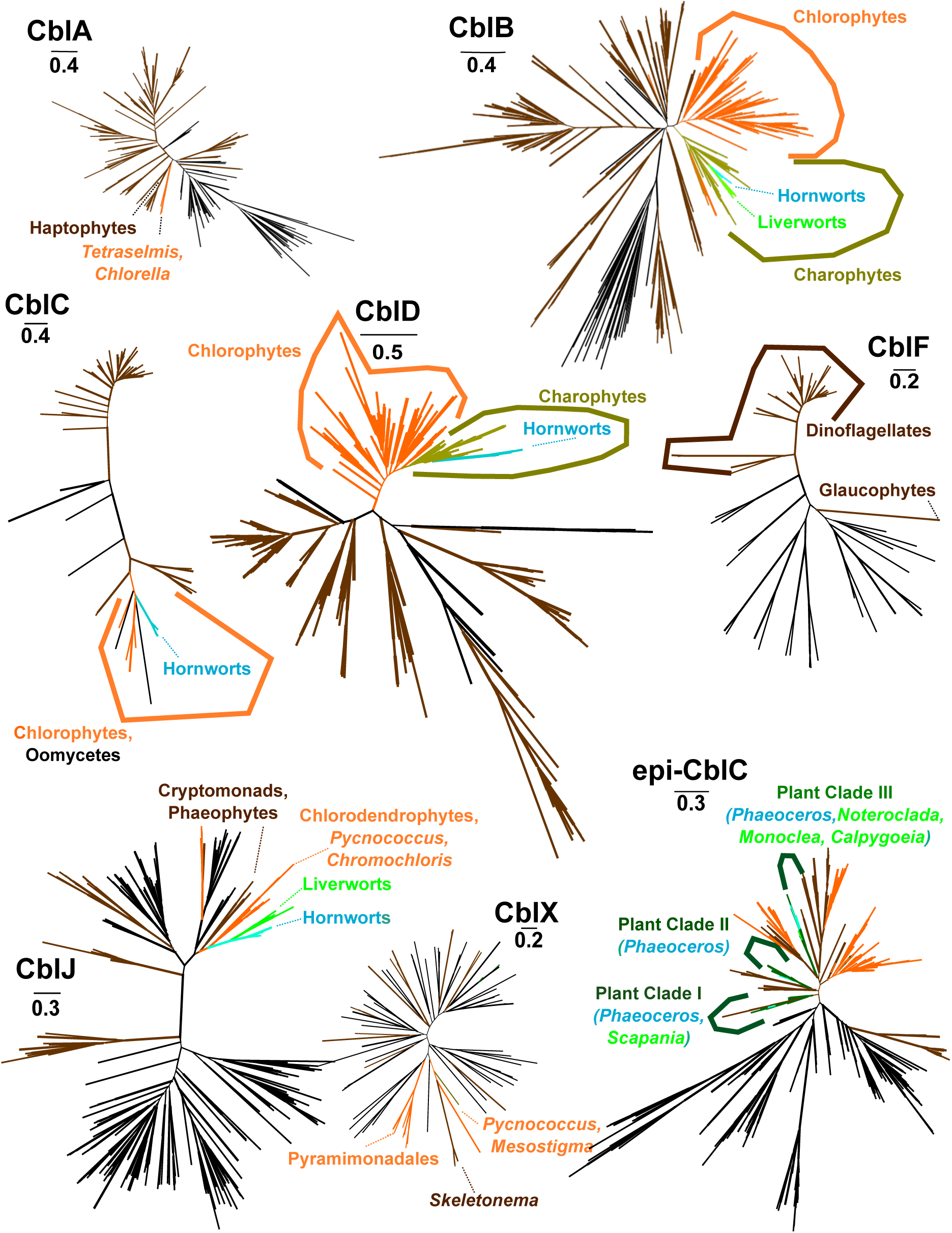
Phylogenetic trees of intracellular transport pathways B_12_ transport across photosynthetic eukaryotes. This figure shows best-scoring RAxML trees realised across 350 bootstrap replicates with the PROTGAMMAJTT substitution matrix for CBLA, CBLB, CBLC, CBLD, CBLF, CBLJ, CBLX and epi-CBLC proteins. Taxa are phylogenetic origin: dark green-vascular plants and mosses; light green-liverworts; cyan-hornworts; dark yellow-other streptophytes; orange-chlorophytes; dark brown-other algae; black-other lineages. Branch thickness corresponds to bootstrap support.

The only protein for which we could identify consistent endomembrane and/ or vacuolar targeting predictions amongst our recovered homologues was for CblF (**Fig. S2**), but this was almost exclusively detected amongst dinoflagellates in our dataset (**Figs. 2, 4**). We therefore tentatively propose that the B_12_ uptake and intracellular trafficking pathways may not be identical across different branches of the eukaryotic tree of life. We could detect potential homologues of both CblX and epi-CblC in hornworts, but found these also in vascular plants and with no clear underlying phylogenetic signal, so we are not confident of the function of these enzymes in B_12_-associated hornwort metabolism (**Figs. 2, 4**).

Surprisingly, homologues of CblB, an enzyme responsible for the synthesis of adenosyl-cobalamin [7], were found in hornworts (**Figs. 2, S1**). This was despite the absence of MCM, which typically uses adenosyl-cobalamin, alongside the CblB partner enzyme CblA from the streptophytes (**Fig. 2**) [31]. We could further detect homologues of CblB and indeed CblJ in liverworts, and in both cases the homologues from these lineages grouped monophyletically with hornworts (**Fig. 4**) and possess well-conserved PFAM domains (**Fig. S1**). It is therefore possible that some form of adenosyl-cobalamin dependent metabolism persisted in the last common plant ancestor, and that this metabolism is retained not only by hornworts but by liverworts within the bryophyte lineage. The validation of this pathway will depend on the functional characterisation of putative adenosyl-cobalamin dependent enzymes, e.g., from published liverwort and hornwort genomes [44, 74].

### Rare occurrences of B_12_-independent algae are biased towards terrestrial and symbiotic species

Finally, we considered across our global datasets which photosynthetic eukaryotes, other than plants, may have secondarily lost B_12_-dependent metabolism completely. Considering B_12_ independence as the presence of METE but absence of METH/ MTRR, RNR II or MCM activities, we identify seven phylogenetically validated occurrences of the loss of B_12_-associated metabolism (**Table 2**). For four species, the same distributions could be identified from PFAM domains only. Finally, we identified four species that possess METE and either MCM or RNR II, but lack METH/MTRR activities, suggesting a loss of B_12_-related methionine synthesis (**Table 2**).

The widespread but seemingly random loss of METE led us to consider what environmental factors might be involved in driving the distribution of this trait, which might be considered *a priori* deleterious. An interactive map of all collection sites reported for our dataset is directly available at https://tinyurl.com/3esfpkmv. The B_12_-independent species in our dataset (blue points on interactive map, and listed in **Table 2**) show characteristic distributions considering their collection sites, mainly originating from freshwater (e.g., *Leptosira obovata, Porphyridium cruentum, Poteriospumella* JBC07) or terrestrial habitats (*Coccomyxa pringsheimii, Cyanidioschyzon merolae, Galdieria sulphuraria*). B_12_-producing bacteria are abundant in some freshwater habitats (e.g., in sediments, as fish gut commensals) but appear to be present in much lower quantities in the water column (e.g., 10^3^-fold less than in sediments), calling into question the bioavailability of B_12_ to freshwater algae [75, 76]. It is possible in some habitats that this is due to cobalt scarcity [77], but is not universally the case. We similarly note that *Geminigera cryophila*, which not only lacks METH but is also the only cryptomonad within our dataset to encode METE, was collected from Antarctic water masses, which have been previously shown to be characterised by vitamin B_12_-limitation and co-limitation (**Fig. 1**) [78, 79]. Finally, many of the B_12_-independent algal species are endobiotic to other organisms (*Leptosira obovata* and two members of the order Trentepohliales, isolated from lichens [80]; *Vitrella brassicaformis*, a coral symbiont [68]; and *Brandtodinium nutriculum*, a foraminiferan symbiont [81]), and may be subject to chronic B_12_ limitation in their native habitats, or even potentially receive methionine and other key metabolites such as folate directly from their hosts.

## Discussion

In this study, we query the distribution of thirteen B_12_-associated proteins across more than 1,600 genomes and transcriptomes from plastid-containing eukaryotes (plants and eukaryotic algae), in particular profiting from the substantial advance in plant genomics made through the OneKp transcriptome project [38] (**Fig. 1**, **Table 2**). Our data suggest the hitherto undocumented retention of vertically inherited B_12_-associated methionine synthesis pathways (METH, MTRR **Figs. 2, 3**) and B_12_ uptake-associated proteins (CblB, C D, J ; **Figs. 2, 4**) in hornworts, and the further retention of potential CblB and CblJ homologues in liverworts (**Fig. 4**). Formally, the active use of B_12_ by hornworts awaits functional characterisation. This may be best performed by bioassay of B_12_ content in hornwort tissue [23], but might also come from the experimental inference of B_12_ use, via B_12_-dependent downregulation of hornwort METE expression [37], or the uptake of fluorescently-labelled B_12_ analogues [82] by hornwort cells.

Considering the predicted functionality and vertical origin of the hornwort homologues, our data indicate the probable presence of B_12_-dependent metabolism in the last common plant ancestor following the transition to land, with subsequent losses in mosses, vascular plants, and (dependent on the functions of the detected CblB and CblJ homologues), liverworts (**Fig. 5**). It remains to be determined at what point B_12_-dependent metabolism was lost during land plant evolution. Some phylogenetic studies have proposed hornworts as the earliest-diverging land plant group [83, 84], which would imply a single loss of METH in a common ancestor of liverworts, mosses and vascular plants. Other studies, including the multigene phylogenomic analyses performed as part of the *Anthoceros* genome and OneKp transcriptome projects [38, 43, 44], recover well-supported monophyly of bryophytes, to the exclusion of vascular plants (**Fig. 5**). This topology would indicate independent losses of B_12_-dependent metabolism in vascular plants and mosses, with a potential third loss of use of methylcobalamin, but retention of MCM and its requirement for adenosyl-cobalamin, in liverworts (**Fig. 5**). In either case, the presence of a metabolic activity in hornworts previously considered unique to algae further blurs the biological distinctions between aquatic streptophytes and plants. This is alongside the retention of biophysical carbon concentrating mechanisms in hornworts [55, 85], and the preceding evolution of plant-like auxin transporters and homeobox transcription factors in charophytes [41, 42].

**Fig. 5.**
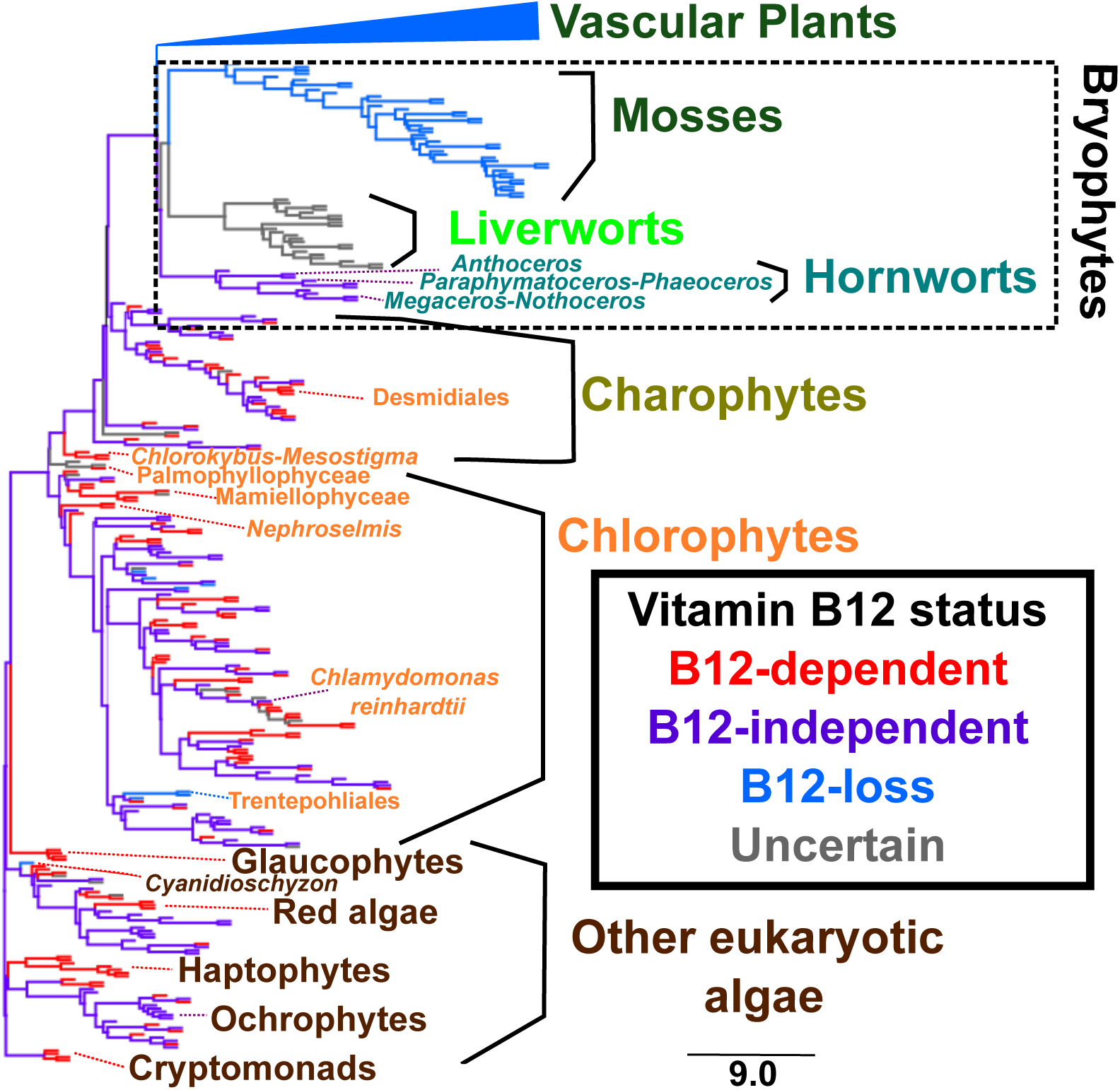
Vertical inheritance of hornwort B_12_ metabolism. This figure shows a concatenated multigene tree topology for all OneKp transcriptomes, taken from [38]. Taxa names are shown by taxonomic affiliation: dark green-vascular plants and mosses; light green-liverworts; cyan-hornworts; dark yellow-other streptophytes; orange-chlorophytes; dark brown-other algae. Branches are shaded by inferred B_12_ status: red-presence of B_12_-dependent metabolism only; purple-presence of B_12_-dependent and –independent metabolism; light blue: B_12_-independent metabolism only; grey-unknown B_12_ status. For concision, vascular plants (universally lacking B_12_-dependent metabolism) are collapsed to a single branch. Liverworts are marked as « unknown » due to the uncertain function of their encoded CblB and CblJ proteins. Major algal lineages are labelled, alongside invididual species of interest and algal clades with inferred losses of B_12_-associated metabolism. ; and a nexus format of the schematic OneKp concatenated topology in **Dataset S1**, sheet 21.

Finally, it remains to be determined why B_12_-dependent metabolism was lost in early plant lineages. Many hornwort species are characterised by the presence of cyanobacterial symbionts, which might produce pseudocobalamin; this is not bioavailable to microalgal species [27, 86]. It remains to be determined whether hornworts can assimilate pseudocobalamin and/ or remodel it into cobalamin, and if this is more bioavailable to them than other plant lineages from which METH has been lost. Or indeed if the plant microbiome has played a broader role in the loss of cobalamin-dependent metabolism. Considering the widespread distribution of B_12_-independent metabolism across the algal tree of life (**Table 2**), we note repeated losses of METH and its associated enzymes in other freshwater, terrestrial and symbiotic lineages. Thus comparative genomic and physiological studies of these species may elucidate the ecological reasons for the loss of B_12_-associated metabolism both in algae and land plants. Understanding why plants do not utilise B_12_-dependent enzymes is particularly important due to the prevalence of vitamin B_12_ insufficiency/deficiency in populations consuming plant-based diets [9]. Ultimately, understanding the significance of B_12_ utilisation and acquisition across algal biology may faciliate the synthetic reintroduction of B_12_ uptake into transformable crop species, or the mass cultivation of algae for dietary consumption [82, 87], sustainably feeding the human planetary population [13].

## Materials and Methods

### Homologue detection

Potential homologues of thirteen Vitamin B_12_-associated enzymes (*Chlamydomonas* METE with GBID : AAC49178.1, METH : XP_042923308.1; human CblA : NP_001362573.1, CblB : NP_443077.1, CblC : NP_056321.2, CblD : NP_056517.1, CblF : NP_060838.3, CblJ : NP_001340531.1, CblX : NP_005325.2, epi-CblC : NP_001189360.1, MTRR : AAF16876.1 ; *Phaeodactylum* MCM : XP_002179504.1 ; *Euglena* RNR II : Q2PDF6.1) [36] were searched across a composite library of 1,663 non-redundant plant and algal genomes and transcriptomes (**Table 2**) [38, 47, 49, 50] by BLASTp with threshold e-value 10^-05^ [88]. Potential matching sequences were extracted and searched by BLASTp against the complete *Arabidopsis thaliana* genome [89], which uniquely encodes METE, supplemented with the query sequences defined above. Sequences that retrieved a best-scoring match to a query sequence were retained for downstream phylogenetic analysis (**Dataset S1**). For *METE* which is retained in b-plants, genes that retrieved one of the three *Arabidopsis* methionine synthases (ATMS1-At5g17920; ATMS2 At3g03780; ATMS3-At5g20980) were likewise retained for downstream analysis [58, 90].

### Phylogeny

Inferred homologues were aligned against the query sequence, and best-scoring homologues obtained from parallel BLASTp searches of 51 combined genome and transcriptome libraries corresponding to different prokaryotic and non-photosynthetic eukaryotic taxonomic groups from across the tree of life [49] by MAFFT v 7.487 using the –gt auto setting [91]. The resulting alignments were imported into GeneIOUS v 10.0.9 and initially screened using the in-built NJ tree function with 100 replicates and random starting seeds; and highly divergent branches (defined visually as branches with > 1.0 calculated substitutions/ site) were iteratively removed [92]. The curated alignment was then trimmed with trimal v 1.4 using the –gt 0.5 setting, prior to a second round of tree building and manual curation [93]. Curated trimmed alignments were then inspected with RAxML v 8.0 using the PROTGAMMAJTT subsitution matrix and 350 bootstrap replicates, following the methodology of previous studies [49]; and the best-scoring tree was then inspected for branches of contaminant origin (defined as sequences from one single library that resolved within or as a sister-group to a distantly related clade, e.g. algal sequences that resolved with bacterial homologues) [51]. The full length sequences of homologues that passed the initial RAxML phylogenetic curation were finally passed through a second iteration of mafft alignment, manual curation, and RAxML phylogeny, and confirmed to resolve in topologies coherent with protein function and taxonomy prior to enumeration of homologue presence/ absence (**Dataset S1**).

### Functional, targeting and biogeographical annotation

PFAM domains were searched in each homologue retrieved by the initial RbH search by HMMER v 3.3.2 against the Pfam v 35.0 library [52, 94]. Sequences that retrieved PFAM domains associated with each query protein were recorded (METE : Meth_synt_1- PF08267, Meth_synt_2- PF01717; METH : B_12_-binding- PF02310, B_12_-binding_2- PF02607, Met_synt_B_12_- PF02965, Pterin_bind- PF00809, S-methyl_trans- PF02574 ; CblA : MeaB- PF03308 ; CblB : Cob_adeno_trans- PF01923 ; CblC : MMACHC- PF16690 ; CblD : MMADHC- PF10229 ; CblF : LMBR1- PF04791 ; CblJ : ABC_tran- PF00005 ; CblX : Kelch_1- PF01344, Kelch_2- PF07646, Kelch_3- PF13415, Kelch_4- PF13418, Kelch_5- PF13854, Kelch_6- PF13964 ; epi-CblC : AhpC_TSA- PF00578, Redoxin- PF08534, 1-cysPrx_C- PF10417MCM : MM_CoA_mutase- PF01642 ; MTRR : Flavodoxin_1- PF00258, FAD_binding_1- PF00667, NAD_binding_1- PF00175 ; RNR II- Ribonuc_red_lgC- PF02867, RNR II_Alpha- PF17975) with threshold e-value 10^-05^. Localisations for each protein were inferred, considering both the full length of the protein sequence and the protein sequence trimmed to the first annotated methionine, using WolfPSort v 1.0 with plant substitution matrix [95]; SignalP v 5.0 under eukaryotic settings [96]; TargetP v 2.0 with plant substitution matrix [97]; and HECTAR v 1.0 under default settings [98]. Collection sites were recorded for each algal species using the corresponding culture collection records considering strain synonyms, and where appropriate direct information from the literature or collector [99–107]. PFAM boxplots in **Fig. S1** are displayed using BoxPlotR [108].

### Data deposition

All strain information, recovered homologues by RbH, alignments, phylogenetic topologies, PFAM and targeting annotations are provided in **Dataset S1**. An interactive map of all algal collection sites identified within the dataset, shadeable either via phylogenetic affiliation or inferred Vitamin B_12_ metabolic status, is available via https://tinyurl.com/3esfpkmv.

### Funding information

RGD acknowledges an ERC Starting Grant (« ChloroMosaic » ; grant number 101039760), awarded 2023-2027. Work at the IBENS is supported by the Investissements d’Avenir programmes PSL and MEMOLIFE: MEMO LIFE (ANR-10-LABX-54), and PSL*Research University (ANR-11-IDEX-0001-02).

## Acknowledgments

The authors thank Dr. Fabio Rocha Jimenez Vieira (IBENS) for assistance with the decontamination of the OneKp transcriptome dataset; and Ms. Cathy Johnston (CSIRO), Dr. Maike Lorenz (Georg-August-Universität Göttingen), Prof. Michael Melkonian (Universität Köln) and Mr. Stephen Peña (UTEX Culture Collection) for the assistance of strain isolation sites tabulated in **Dataset S1**.

---

**Fig. S1. Metabolic functions of Vitamin B_12_-associated metabolism in plants.** This figure shows boxplots, displayed using BoxPlotR [108], of the –log_10_ of the calculated hmmer e-value scores of key PFAM domains observed for each protein [94] of all phylogenetically reconciled homologues of eight potential B_12_-associated proteins found in hornworts : METE, METH, MTRR, CBLB, CBLC, CBLD, CBLJ and epi-CBLC. Values are shown for four taxonomic groups: eukaryotic algae ; hornworts ; liverworts ; and mosses and vascular plants (grouped together by the absence of B_12_ metabolism as opposed to phylogeny). Greater values imply greater fidelity to the PFAM hmm ; groups for which no homologues were identified are plotted as a −1 value. For the METH sequences, an alignment is additionally provided of a 20 aa region of the first substrate binding pocket of the pterin binding domain of hornwort METH against the equivalent *Chlamydomonas* sequence following [21], and showing broad conservation of key residues and inferred functions. An overview of the PFAM values is given in **Dataset S1**, sheet 3; with individual scores for each homologue provided in **Dataset S1**, sheets 5-17.

**Fig. S2. Targeting predictions of individual B_12_-associated proteins. A** : bar plots of the proportion of phylogenetically identified homologues of each B_12_-associated protein to possess at least one mitochondrial, chloroplast or endomembrane targeting sequence, from combined WolfPSort, HECTAR, SignalP and TargetP predictions. Except for MCM which shows broadly mitochondrial targeting predictions and CblF with predominantly endomembrane localisations, no patterns are consistently observed. **B** : wheel plots of the taxonomic composition of mitochondria-targeted MCM sequences, showing broad conservation of this localisation across algae with secondary red chloroplasts ; and of endomembrane-targeted CblF sequences, which are restricted to dinoflagellates.

## Notes

https://tinyurl.com/3esfpkmv

### Competing Interest Statement

The authors have declared no competing interest.

