## Supplementary material for "Presence of vitamin B_12_ metabolism in the last common ancestor of land plants": Fig. S1

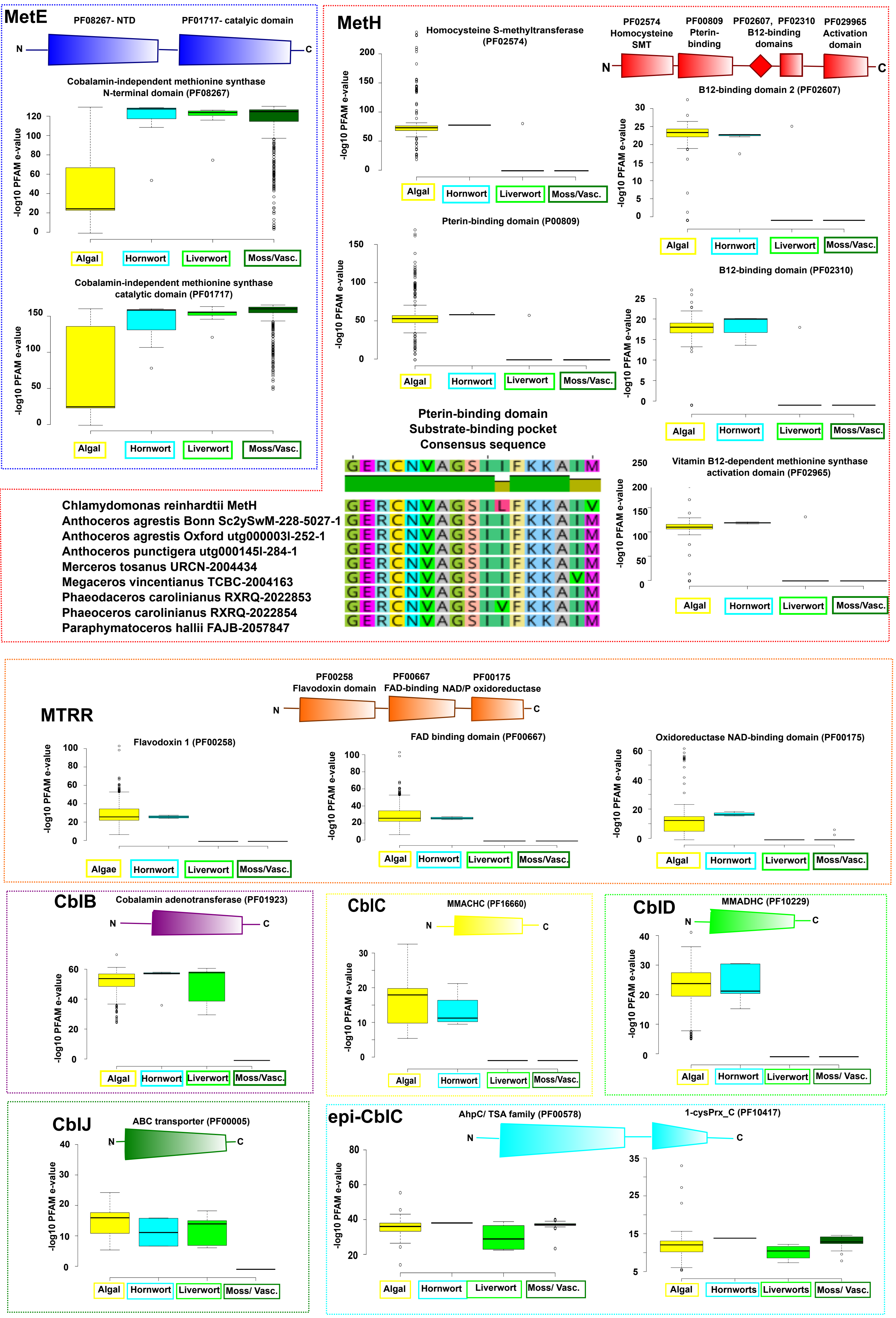


**Fig. S1**. This figure shows boxplots, displayed using BoxPlotR, of the -log10 of the calculated hmmer e-value scores of key PFAM domains observed for each protein, of all phylogenetically reconciled homologues of eight potential B12-associated proteins found in hornworts: METE, METH, MTRR, CBLB, CBLC, CBLD, CBLJ and epi-CBLC. Values are shown for four taxonomic groups: eukaryotic algae; hornworts; liverworts; and mosses and vascular plants (grouped together by the absence of B12-metabolism as opposed to phylogeny). Greater values imply greater fidelity to the PFAM hmm; groups for which no homologues were identified are plotted as a -1 value. For the METH sequences, an alignment is additionally provided of a 20 aa region of the first substrate binding pocket of the pterin binding domain of hornwort METH against the equivalent Chlamydomonas sequence, and showing broad conservation of key residues and inferred functions. An overview of the PFAM values is given in Dataset S1, sheet 3; with inidividual scores for each homologue provided in Dataset S1, sheet 5-17.
