## Supplementary material for "Presence of vitamin B_12_ metabolism in the last common ancestor of land plants": Fig. S2.

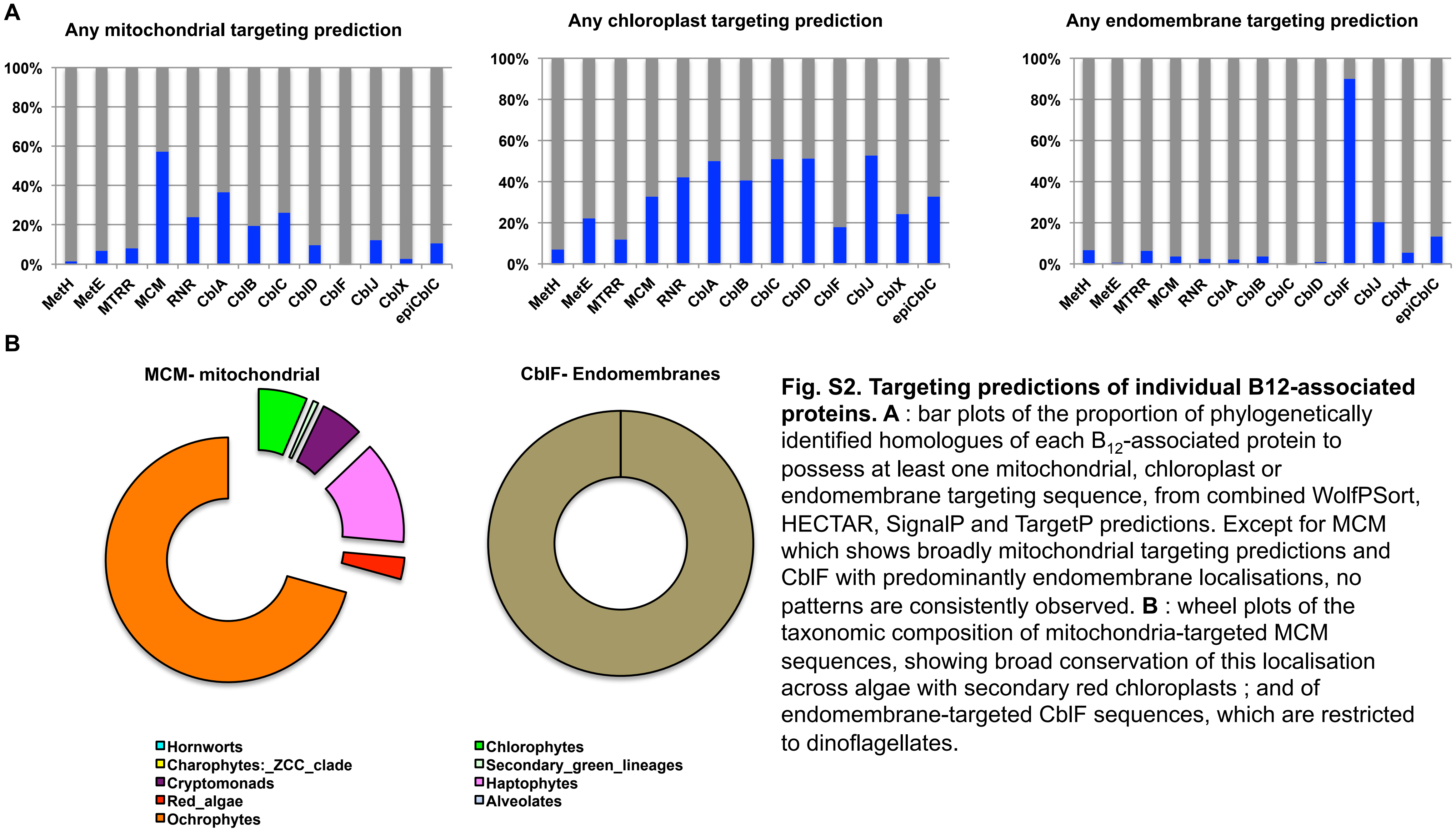


**Fig. S2. Targeting predictions of individual B12-associated proteins. A** : bar plots of the proportion of phylogenetically identified homologues of each $B_{12}$-associated protein to possess at least one mitochondrial, chloroplast or endomembrane targeting sequence, from combined WolfPSort, HECTAR, SignalP and TargetP predictions. Except for MCM which shows broadly mitochondrial targeting predictions and CblF with predominantly endomembrane localisations, no patterns are consistently observed. **B** : wheel plots of the taxonomic composition of mitochondria-targeted MCM sequences, showing broad conservation of this localisation across algae with secondary red chloroplasts ; and of endomembrane-targeted CblF sequences, which are restricted to dinoflagellates.
